# Perinatal development of structural thalamocortical connectivity

**DOI:** 10.1101/2024.04.29.591770

**Authors:** Stuart Oldham, Sina Mansour L., Gareth Ball

**Affiliations:** Developmental Imaging, Murdoch Children’s Research Institute, The Royal Children’s Hospital Melbourne; The Turner Institute for Brain and Mental Health, School of Psychological Sciences and Monash Biomedical Imaging, Monash University, Clayton, Australia; Systems Lab, Department of Psychiatry, The University of Melbourne, Parkville, Victoria, Australia; Centre for Sleep & Cognition & Centre for Translational Magnetic Resonance Research, Yong Loo Lin School of Medicine, National University of Singapore; Department of Paediatrics, University of Melbourne, Melbourne

**Keywords:** thalamus, gradients, perinatal, neurodevelopment, thalamocortical, connectivity

## Abstract

Thalamocortical connections are crucial for relaying sensory information in the brain and facilitate essential functions including motor skills, emotion, and cognition. Emerging evidence suggests that thalamocortical connections are organised along spatial gradients that may reflect their sequential formation during early brain development. However, this has not been extensively characterised in humans. To examine early thalamocortical development, we analysed diffusion MRI data from 345 infants, scanned between 29-45 weeks gestational age. Using diffusion tractography, we mapped thalamocortical connectivity in each neonate and used Principal Component Analysis to extract shared spatial patterns of connectivity. We identified a primary axis of connectivity that varied along an anterior/medial to posterior/lateral gradient within the thalamus, with corresponding projections to cortical areas varying along a rostral-caudal direction. The primary patterns of thalamocortical connectivity were present at 30 weeks’ gestational age and gradually refined during gestation. This refinement was largely driven by the maturation of connections between the thalamus and cortical association areas. Differences in thalamocortical connectivity between preterm and term neonates were only weakly related to primary thalamocortical gradients, suggesting a relative preservation of these features following premature birth. Overall, our results indicate that the organisation of structural thalamocortical connections are highly conserved across individuals, develop early in gestation and gradually mature with age.

## 1 Introduction

The connections between the thalamus and cortex are essential for maintaining sensory and motor control, in addition to higher order functions including attention, memory, emotion, and consciousness (Jankowski et al., 2013; Mitchell, 2015; Sherman, 2016; Sherman & Guillery, 2002; Sommer, 2003). It is widely considered that the diverse functionality of thalamus is underpinned by its nuclear organisation – consisting of 50-60 distinct nuclei with distinctive patterns of molecular, cytoarchitectural, and connectivity properties (Fama & Sullivan, 2015; Jones, 2007; Morel, 2007).

Recent evidence suggests that, in addition to a well characterised nuclear structure, the thalamus exhibits continuous variations in patterns of connectivity, cytoarchitecture, and molecular identity that extend both within and across specific nuclei (Gao et al., 2020; Howell et al., 2024; John et al., 2024; Jones, 1998; Li et al., 2020; Mai & Majtanik, 2019; McFarland & Haber, 2002; Oldham & Ball, 2023; Park et al., 2024; Phillips et al., 2019; Roy et al., 2022; Saunders et al., 2018). The spatial organisation of the thalamus along continuous axes is reflected by concerted variations in gene transcription, axonal morphology, laminar targeting, and electrophysiological properties, alongside other key principles of cortical organisation (Oldham & Ball, 2023; Phillips et al., 2019).

The presence of continuous macroscopic variation in thalamic connectivity presents evidence that thalamocortical patterning is shaped by morphogenetic gradients during early brain development (Govek et al., 2022; Nagalski et al., 2016; Price et al., 2012; Teissier & Pierani, 2021). Firstly, the orientation of the principal axes of variation correspond to known developmental gradients (Altman & Bayer, 1979; Cahalane et al., 2012; Finlay & Uchiyama, 2015; Nakagawa, 2019; Wong et al., 2018). Secondly, genes expressed along the primary molecular and connectomic axes are differentially expressed during the prenatal period (Oldham & Ball, 2023). Finally, these genes are enriched for neurodevelopmental disorders such as schizophrenia (Elvsåshagen et al., 2021; Oldham & Ball, 2023).

Despite its central importance in brain organisation and function, only a relatively limited number of neuroimaging studies have addressed the development of thalamocortical connectivity in the human brain during the perinatal period (Ball et al., 2012, 2013, 2015; Batalle et al., 2017; Jakab et al., 2020; Sa De Almeida et al., 2021; Toulmin et al., 2021; Wilson et al., 2023; Zheng et al., 2023). The second half of gestation is a critical period for the formation of thalamocortical connections (Kostović et al., 2019, 2021; Kostović & Judaš, 2010). By birth, the overall patterning of structural thalamocortical connections is established (Kostović et al., 2019), with maturation progressing from early-maturing primary sensory areas to later-maturing association cortex in the frontal lobe (Zheng et al., 2023). Evidence suggests that interruptions to early brain development due to preterm birth can negatively impact the structural, and subsequently functional, connectivity of the thalamus in infancy, leading to poor cognitive and motor outcomes (Alcauter et al., 2014; Ball et al., 2012, 2015; Jakab et al., 2020; Toulmin et al., 2015, 2021). Investigating the spatial and temporal progression in how structural thalamocortical connections form and mature is vital to understanding the development of healthy and abnormal brain function.

While other studies have outlined the broad developmental trends of structural thalamocortical connectivity, previous examples have either examined a limited number of major thalamocortical tracts or only used a coarse cortical parcellation to define cortical connectivity. In this study, we employ a high-resolution connectome approach to estimate dense maps of thalamocortical structural connectivity from 29 to 45 weeks gestation, examining the development of organisational thalamocortical gradients and providing a granular analysis of early thalamic structural connectivity to the cortex. In addition, we test the hypothesis that preterm birth negatively impacts early patterning of structural thalamocortical connectivity.

## 2 Methods

Participant data was acquired from the third release of the Developing Human Connectome Project (dHCP; ethics approved by the United Kingdom Health Research Ethics Authority, reference no. 14/LO/1169) (Edwards et al., 2022; Hughes et al., 2017) which consisted of 783 neonates (360 female; median birth age [range] = 39^+2^ weeks [23-43^+4^]) across 889 scans (median scan age [range] = 40^+6^ [26^+5^-45^+1^] weeks; 107 neonates were scanned multiple times). Following strict quality control (see below), 363 scans were retained for analysis. For the 18 neonates with multiple scans, we selected the scan closest to their birth age. The final cohort comprised 345 neonates (165 females; median birth age [range] = 39 weeks [23^+4^- 42^+2^]; median scan age [range] = 40^+3^ [29^+2^-45^+1^] weeks). Of these, we selected the oldest 20 by scan age, who were born at term age and had a radiological score of 1 (indicating no radiological abnormalities or pathologies), to create a term template which acted as a reference.

### 2.1 MRI acquisition and processing

Images were acquired on a Phillips Achieva 3T scanner at St Thomas Hospital, London, United Kingdom (Hughes et al., 2017). T2-weighted images were acquired using multislice fast spin-echo sequence with TR = 12000 ms, TE = 156 ms, using overlapping slices (0.8 × 0.8 × 1.6 mm). Diffusion data was acquired with TR = 4000ms, TE = 90ms, 20 b=0s/mm^2^ volumes and 64 400s/mm^2^, 88 1000s/mm^2^ and 128 2600s/mm^2^ b-value volumes, and 1.5×1.5×3mm voxels in 64 slices.

Structural images were processed using the dHCP’s minimal preprocessing pipeline (Makropoulos et al., 2018). This included applying bias correction and brain extraction. The DRAW-EM algorithm was then used to create tissue segmentations based on the T2- weighted images.

A neonatal-specific processing pipeline was applied to the diffusion data, the full details of which are described elsewhere (Bastiani et al., 2019). In brief, this involved selecting b0 volumes least affected by within-volume motion and using this to estimate the off-response field using FSL’s TOPUP (Andersson et al., 2003) followed by distortion corrections using FSL’s EDDY (Andersson et al., 2016, 2017, 2018; Andersson & Sotiropoulos, 2016). A super-resolution algorithm was applied to achieve an isotropic resolution of 1.5 mm (Kuklisova-Murgasova et al., 2012). The diffusion data was aligned to the individual T2w images (Greve & Fischl, 2009; Jenkinson et al., 2002), and then to the 40-week neonatal dHCP template (Schuh et al., 2018) using nonlinear registration (Andersson et al., 2010).

To ensure only high-quality scans without major motion-related artefacts were used, we examined the quality control summaries of the dHCP diffusion processing pipeline. Scans which were more than two standard deviations away from the mean on any of the volume-to-volume motion, within-volume motion, susceptibility-induced distortions, and eddy current-induced distortions metrics were excluded (n=626).

### 2.2 Thalamic seed definition

Using the thalamic mask manually defined on the dHCP extended 40-week template (Bozek et al., 2018), we placed 800 seeds (arranged in a 1.75mm 3D grid) evenly distributed throughout the thalamic volume in the left hemisphere (**Figure 1A**). Each seed was transformed from the template to each participant’s diffusion scan using the pre-calculated non-linear transformations. Thalamic seed registrations were inspected visually to ensure correct alignment.

**Figure 1.**
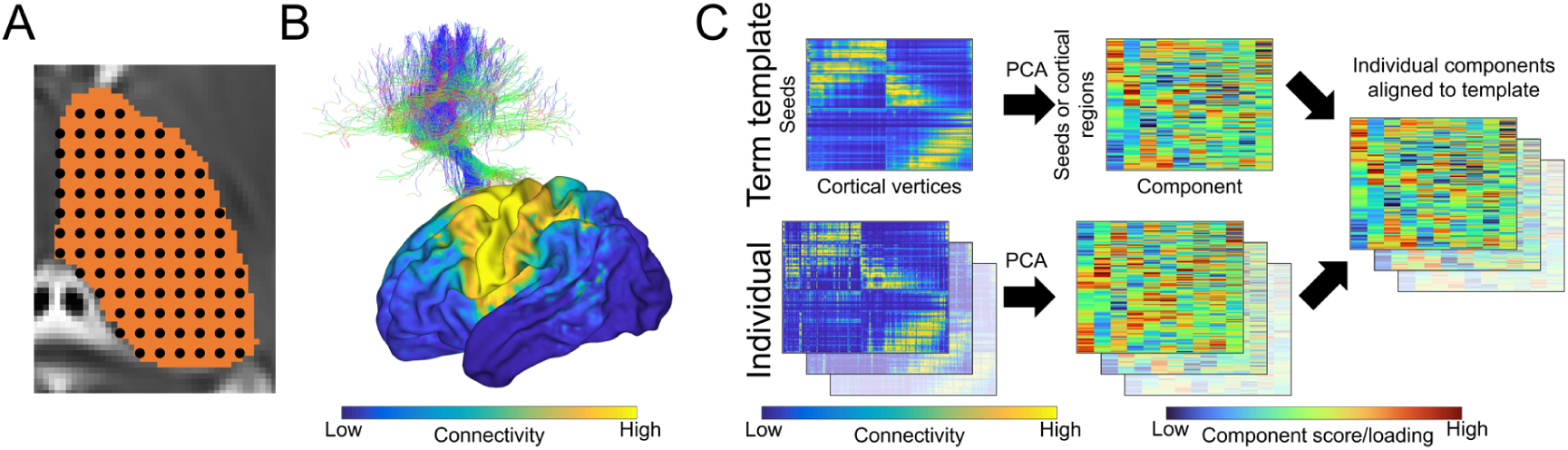
Schematic of processing steps. (**A**) 800 seeds are defined across the thalamic volume and are registered to each individual’s diffusion data. (**B**) Connectivity between each seed and 28,766 cortical vertices is estimated with probabilistic tractography (example connectivity from one thalamic seed is shown; top), producing a dense thalamic-seed-by-cortical-vertex connectivity matrix. (**C**) A term connectome template was created by averaging the connectivity of the oldest 20 term-born neonates, followed by a linear decomposition to construct principal thalamocortical gradients. The remaining neonate’s connectivity matrices were then individually decomposed and the resulting decompositions were aligned to the template gradients for comparison.

### 2.3 Thalamic seed connectivity

For each neonate, we generated 5,000 streamlines from each of the 800 thalamic seeds using MRtrix3 (Jeurissen et al., 2014; Tournier et al., 2004, 2019). To calculate the fibre orientation distributions (FODs) needed for tractography, we first extracted only the 0 and 1000s/mm² b-value volumes for the diffusion data were extracted, as using just these to define FODs using single-shell 3-tissue constrained spherical deconvolution (CSD) has shown better definition of crossing fibres than multi-shell CSD approaches in neonatal data (Dhollander et al., 2019). Next, the response function was estimated using 20 of the oldest term-born subjects to ensure only estimates of relatively mature white matter were used. White matter, grey matter, and cerebral spinal fluid (CSF) response functions were estimated using the *dhollander* algorithm in MRtrix3 (Dhollander et al., 2016). The estimated response functions were used in single-shell 3-tissue CSD to obtain FODs for every participant (Dhollander et al., 2019). A five-tissue-type image was created using segmentations of the grey matter, white matter, and CSF provided by the dHCP to apply Anatomically Constrained Tractography (R. E. Smith et al., 2012). Connectivity between the thalamus and cortex was then calculated using second-order integration over fibre orientation distributions (iFOD2) tractography (Tournier et al., 2010, 2019) (0.75mm step size; 45° maximum angle; 0.05 fibre orientation distribution cut-off). We visually inspected the resulting tractograms to check anatomically plausible streamlines were obtained and no gross abnormalities were present as to ensure our approach was successful in delineating white-matter pathways.

To estimate the spatial distribution of cortical connections from each thalamic seed, we used a surface-based mapping approach. The dHCP neonatal 40-week white matter surface was aligned to each individual’s diffusion space using transforms provided by the dHCP. This provided a surface mesh with matched geometry to each individual’s surface with vertex correspondence across individuals for comparison.

Tractography streamlines were used to map high-resolution thalamocortical connectivity maps (Mansour L et al., 2021). As most thalamocortical connections are ipsilateral (Dermon & Barbas, 1994), we only measured connectivity from the left thalamic seeds to vertices in the left cortical hemisphere. For all thalamic seeds, reconstructed streamlines were assigned to the nearest cortical vertex within a 5mm radius of their endpoint, and each streamline was weighted by the average of sampled mean fractional anisotropy (FA) along its length. The connectivity between each thalamic seed to a given cortical vertex was then taken as the sum of streamline weights. Connectome spatial smoothing (3mm FWHM, 0.01 epsilon) was subsequently performed to account for the susceptibility of high-resolution connectomes to the impacts of accumulated integration errors in streamline propagation (Mansour et al., 2022). This method involves applying a pair of Gaussian smoothing kernels to adjust the strength of connectivity across cortical vertices (**Figure 1B**) and has previously been shown to improve the reliability of individual connectivity measures (Mansour et al., 2022). Using this approach, we created a dense connectome matrix summarising white-matter connectivity between 800 thalamic seeds and all 28,766 cortical vertices of the left hemisphere (excluding those assigned to the medial wall; **Figure 1C**).

Reconstructing the structural connectivity from deep thalamic structures (especially towards medial areas) using tractography is challenging, due to the impact of partial volume effects on these regions. Additionally, a tract seeded from one of these regions needs to traverse many voxels of low anisotropy to reach the cortex, reducing the likelihood that such a connection will be reliably detected by tractography. These difficulties may be compounded by lower signal in neonatal imaging data. We examined average connectivity of each seed to the cortex (across individuals), identifying a set of seeds located near to the midline with very low cortical connectivity (**Figure S1A**). To avoid potential biases from poor tracking from the medial wall, we removed low connectivity seeds (connected to <100 cortical vertices; **Figure S1B**). This resulted in 646 seeds retained for the final analysis (**Figure S1C**).

Cortical connectivity across seeds was normalised using a scaled sigmoid transformation to the interval [0,1]. This first involved applying a sigmoidal transformation to the raw data:

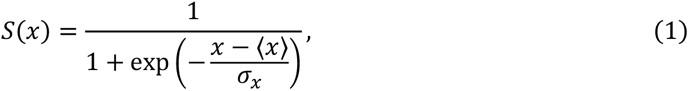

where *S*(*x*) is the normalised value of a connection, *x* is the raw value, ⟨*x*⟩ is the mean and *σ_x_* is the standard deviation of the values of that connection across thalamic seeds. Following the sigmoidal transform, cortical connections were linearly scaled to the unit interval. This transformation was used to reduce the impact of outliers in the data (Fulcher & Fornito, 2016; Parkes et al., 2017).

### 2.4 Gradient decomposition

We decomposed the concatenated 646-by-28766 (*n* × *m*) normalised thalamocortical connectivity matrix *M* (*M* was centred prior to the decomposition) into a set of orthogonal components using Principal Component Analysis (PCA) via Singular Value Decomposition (SVD):

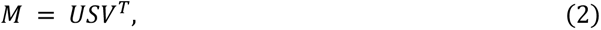

where *U* is a *n* × *n* matrix of left singular vectors; *S* is a *n* × *m* rectangular diagonal matrix of the singular values *s* of *M*; and *V* is a *m* × *m* matrix of right singular vectors. This approach reduces the dimensionality of the data by finding components (patterns of variation which together maximise the variance explained in the data) which are orthogonal to each other. The decomposition is normally truncated to *k* < *min*(*n*, *m*) and the variance explained by each component, λ*_k_*, is given by its singular values, *s_k_*:

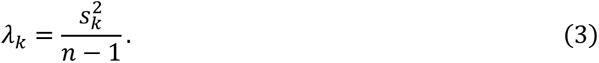

Therefore *US* is a 646 × *k* matrix representing the Principal Component (PC) scores, one per thalamic seed for each of *k* components; and *V* is a 28766 × *k* matrix representing the PC loadings that denote the contributions of each cortical vertex’s (normalised) thalamic connectivity to each component. Each PC indicates a pattern of connectivity that accounts for a proportion of the variance in the data. The loadings for each PC indicate a pattern of cortical connectivity, while the corresponding PC scores indicate how strongly the connectivity from a particular thalamic seed aligns with the cortical loadings. The sign of the score indicates the direction of the association. A positive value means that seed is strongly connected to cortical areas with positive loadings (for that PC), and weakly connected to cortical areas with a negative loadings. Similarly, a negative score indicates the seed is weakly connected to cortical areas with positive loadings, but strongly connected to negative ones. The magnitude of the score indicates the strength of this association.

### 2.5 Gradient alignment

To allow comparison of individual connectivity components across the third trimester, we aligned all PC score matrices using a Procrustes transform. To do so, we selected the 20 oldest neonates who were born at term, averaged their thalamocortical connectivity matrices and decomposed this average matrix via PCA to create a ‘term template’ of scores and loadings. The remaining neonatal thalamocortical connectivity matrices not included in construction of the term template were then individually decomposed using PCA. A Procrustes rotation (based on the first five components as to reduce computational demands) was used to align each individual decomposition to the term decomposition (alignment was performed on the *US* matrix, the resulting transforms derived from this alignment were used to align the *V* matrix; **Figure 1C**).

### 2.6 Cortical null models

Tissue properties (thickness, cytoarchitecture, connectivity) vary smoothly across the cortex with nearby areas sharing similar features, a phenomenon driven by spatial autocorrelation (Markello & Misic, 2021). The presence of spatial autocorrelations can lead to overestimation of correlations between different, smoothly varying cortical properties. To mitigate this effect, we compared spatial correspondences across cortical properties to a spatial-autocorrelation-preserving null model using a permutation test (via spin test) (Alexander-Bloch et al., 2018; Markello & Misic, 2021). The position of cortical vertices was first randomly rotated on the spherical representation of the cortical surface. Each of the rotated vertices is then matched to the closest original vertex, creating a mapping of rotated-to-original vertices which can they be used to rotate the values of a cortical feature. This vertex mapping preserves the spatial autocorrelation and can then be used to conduct permutation testing. We assessed the Pearson correlation between a pair of brain maps and compared it to a distribution of correlations generated from 1,000 null permutations (i.e., one of the brain maps was permuted using the vertex mapping are correlated with the other non-permuted one). This process was then repeated in the opposite direction (i.e., the second brain map was also permuted as to remove any bias from only ever permuting one brain map), and the average *p*-value was taken to obtain a spin-test derived *p*-value (*p*_*spin*_), which was considered significant at < .05 (Alexander-Bloch et al., 2018; Markello & Misic, 2021).

## 3 Results

We used PCA to decompose a matrix of average thalamocortical connectivity from 20 term-born neonates. The first three principal components (PCs) explained 48.6%, 28.6%, and 8.22% of variance, respectively. Thalamic PC scores varied along an anterior/medial to posterior/lateral axis (**Figure 2A**) and were strongly correlated with both medial-lateral (x axis; r = 0.70, *p* < .001; **Figure S2A**) and anterior-posterior seed position (y axis; r = −0.85, *p* < .001; **Figure S2B**), but only weakly correlated with inferior-superior position (z axis; r=-0.27, p<0.001). In contrast, thalamic PC2 scores varied in a radial pattern outward from a lateral anchor point (**Figure 2B**), while PC3 was oriented along an axis varying from anterior/ventral posterolateral areas to posterior/ventral lateral ones (**Figure 2C**). Compared to PC1, the spatial patterns of PC2 and PC3 were only weakly correlated with cardinal image axes (**Figure S2D-i**).

**Figure 2.**
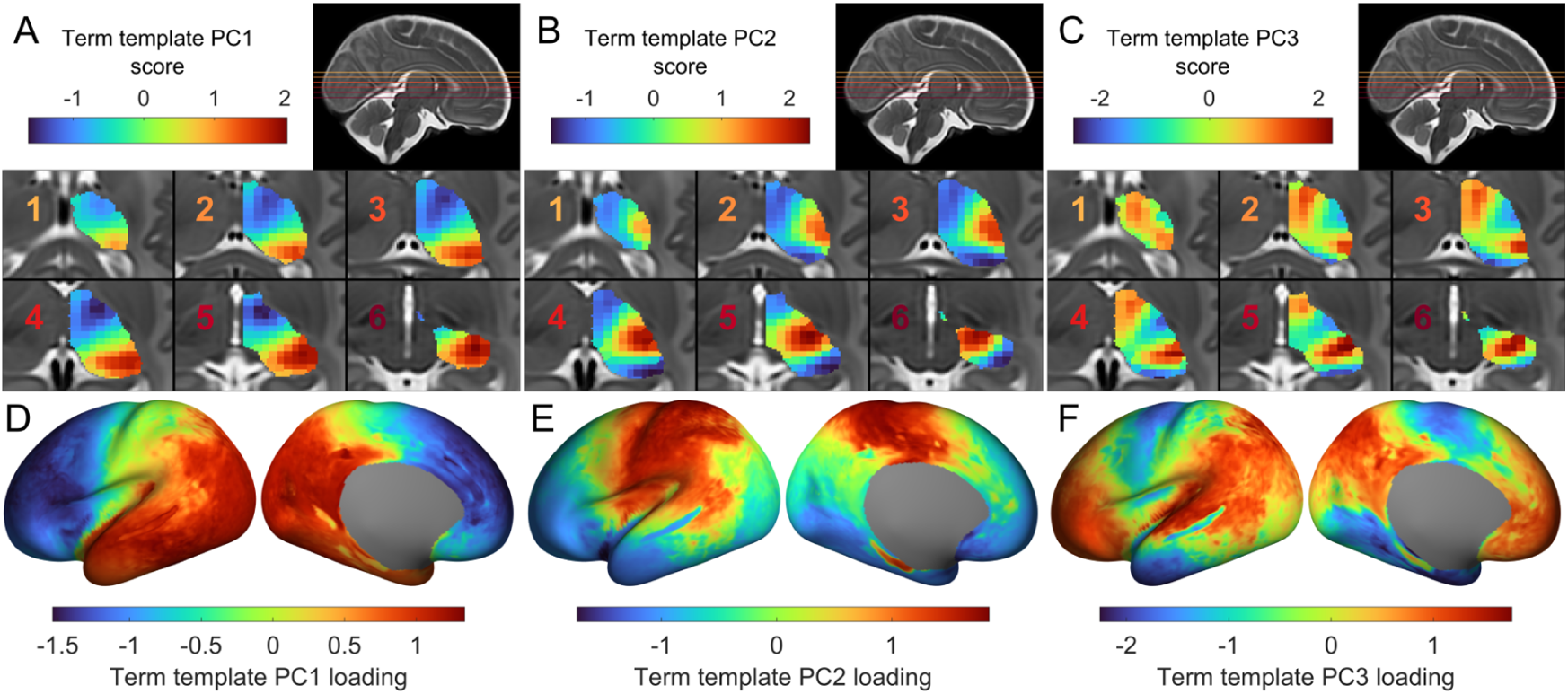
Axes of thalamocortical connectivity. (**A**) Projection of PC1 scores onto thalamic voxels, revealing variation along medial-lateral and anterior-posterior directions. (**B**) Projection of PC2 scores onto thalamic voxels. (**C**) Projection of PC3 scores onto thalamic voxels. PC scores for each seed are projected onto the closest voxels in the thalamic mask, overlaid on six axial sections (inset; the colour of the number corresponds to the slice in the insert). (**D**) The PC1 loadings for cortical regions are shown projected onto the cortical surface, revealing a rostral-caudal gradient of thalamocortical connectivity. (**E**) PC2 loadings for cortical regions projected onto the cortical surface. (**F**) PC3 loadings for cortical regions projected onto the cortical surface. Preferential connectivity between thalamic seeds and cortical vertices are shown by similar colours for the respective PC.

Thalamocortical connectivity is topographically arranged, such that the spatial arrangement of thalamic connections are mirrored in their cortical targets (Jones, 2007). The topography of the thalamic PC scores are, in turn, mirrored by the corresponding PC loadings of each cortical vertex (Oldham & Ball, 2023). Projecting the loadings for PC1 onto their respective cortical vertices revealed a rostral-caudal gradient of connectivity (**Figure 2D**). Rostral cortical areas were negatively loaded, and thus preferentially connected to medial-anterior thalamic regions, while caudal regions were positively loaded, indicating preferential connectivity to thalamic posterior-lateral regions. The cortical loadings for PC2 revealed preferential connectivity between the lateral thalamus and primary sensory and motor cortex along a dorsal-ventral axis (**Figure 2E**), while PC3 revealed preferential connectivity to frontal and parietal association areas (Margulies et al., 2016; Sydnor et al., 2021) (**Figure 2F**).

This analysis demonstrates that a simple, low-dimensional representation of thalamocortical connectivity can efficiently capture macroscale patterns of preferential connectivity between regions of the thalamus and cortex, around the time of birth. We next sought to determine how these patterns are established in the time prior to birth. To measure changes in thalamocortical connectivity across the third trimester, we aligned individual connectivity decompositions from the full cohort (scan age: 29 – 45 weeks) to the average term PC components, the ‘term template’, via Procrustes rotations (Vos De Wael et al., 2020). The amount of variance explained for PC1 (34.4%; SD = 3.2), PC2 (21.5%; SD = 2.1), and PC3 (7.5%; SD =0.9), across individuals was lower than observed in the term PC decomposition, likely reflecting the greater amount of noise present in individual data.

Overall, individual PCA decompositions of thalamocortical connectomes were highly similar to the term template (*r =* 0.90 to 0.99; **Figure 3A**) revealing conserved patterns of connectivity among individuals present from at least 30 weeks gestational age. However, we found that similarity to average term connectome decomposition was significantly correlated with age at time of scan (*r* = 0.76, *p* < .001), with greatest dissimilarity at the earliest timepoints (**Figure 3A**). We confirmed that similar associations were observed between PC loadings across individuals, finding a strong correlation between similarity to the term data and scan age (*r* = 0.78, *p* < .001; **Figure 3B**). Similarity of PC2 and PC3 to the term template data was also significantly associated with age in the thalamus (PC2: *r* = 0.77, *p* < .001, **Figure S3A**; PC3; *r* = 0.77, *p* < .001, **Figure S4A**) and cortex (PC2: *r* = 0.74, *p* < .001, **Figure S3B**; PC3; *r* = 0.80, *p* < .001, **Figure S4B**). We also compared how closely each individual’s decomposition correlated with a template constructed from all other non-duplicate scans, finding highly consistent results (PC1 score: *r* = 0.68, *p* < .001; PC2 loading: r = 0.77, *p* < .001) to those obtained when using the term template.

**Figure 3.**
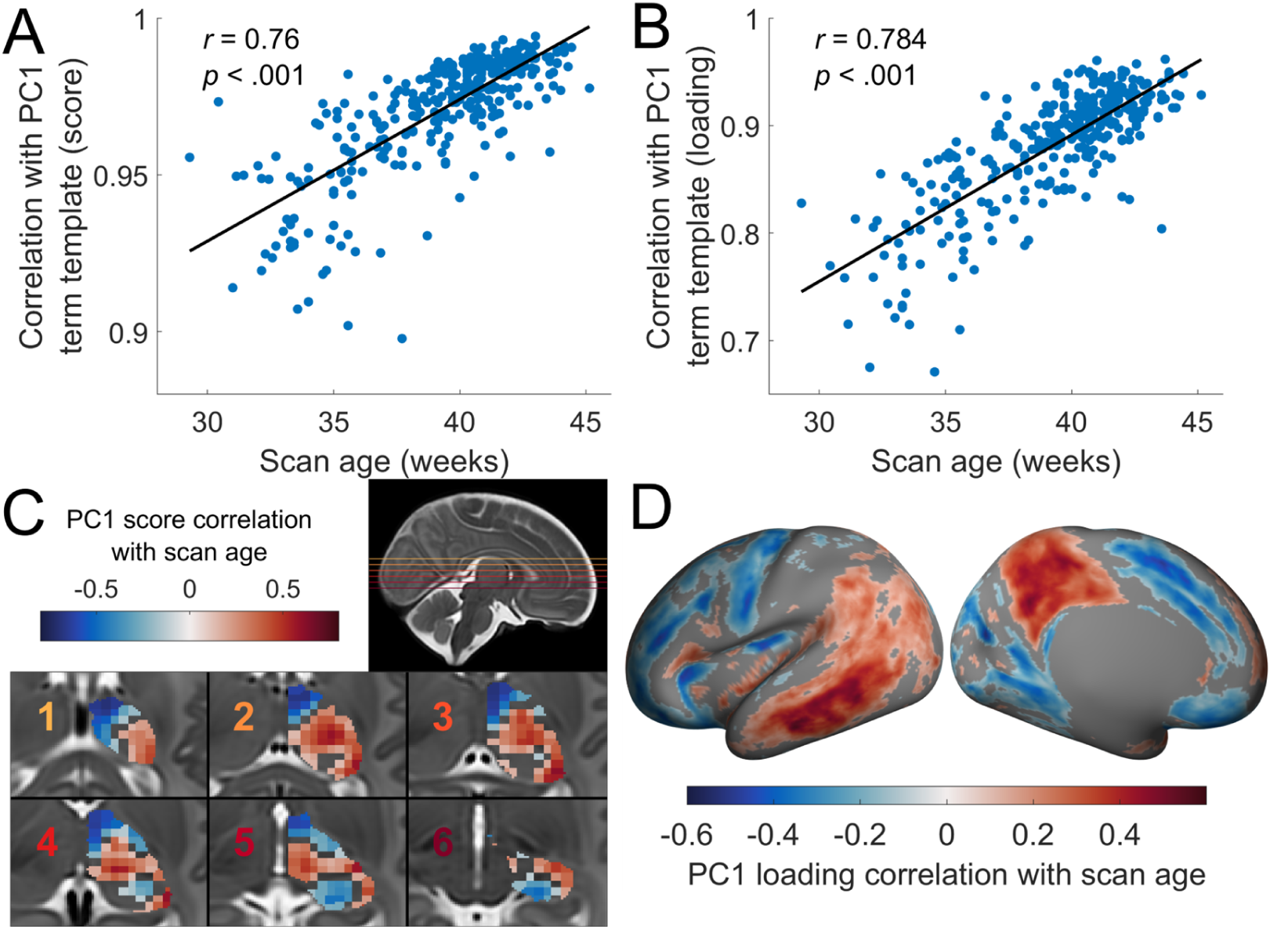
Age-related changes in the primary thalamocortical axis. (**A**) Scatter plot of the correlation between individual and template PC1 thalamic scores and individual scan age. (**B**) Scatter plot of the correlation between individual and template PC1 cortical loadings and individual scan age. (**C**) Correlation between individual PC1 scores and scan age for each thalamic seed (voxels are coloured according to the value of the nearest seed; *p_FDR_* < 0.05; non-significant thalamic areas are not coloured). (**D**) Correlation between individual PC1 loading and scan age for each cortical vertex (*p_FDR_* < 0.05; non-significant vertices are not coloured).

These data suggest a gradual refinement of thalamocortical connectivity towards the time of normal birth. We next sought to establish where in the thalamus and cortex, age-related refinement was most prominent by testing the associations between thalamic seed PC scores and scan age across individuals. As determined by a Pearson correlation with a false-discovery rate correction (Benjamini & Hochberg, 1995) of *p_FDR_* < 0.05, for PC1 we observed decreasing PC scores in the medial-anterior thalamus and pulvinar with increasing age, while ventro-lateral thalamic areas showed increases in PC scores (**Figure 3C**). This reflects an increasing differentiation of thalamic connectivity patterns along PC1. A negative correlation indicates an increasingly negative PC1 score over the third trimester, while a positive correlation indicates an increasingly positive PC1 score. This reflects a shift in the pattern of connectivity between that seed and the cortex over development. For example, the connectivity of seeds with high PC1 score (in the term template) showing a positive correlation with age will have had their connectivity with posterior areas strengthen (relative to anterior areas) across the third trimester. In the cortex, these changes in underlying connectivity are reflected by large age-related changes in PC1 loadings in parietal (increases) and medial frontal areas and motor areas (decreases; **Figure 3D**). Similarly, differentiation between thalamic seed connectivity patterns along PC2 and PC3 become more apparent with age (**Figure S3, S4**).

Age-related changes were only weakly correlated with average term PC1 scores (*r* = 0.13; **Figure S5A)**, suggesting that age-related changes were not simply a strengthening or reinforcement of the existing pattern in the thalamus. A stronger association was observed in cortical vertex PC1 loadings and age-related changes (*r* = 0.44; **Figure S5B**), indicating that areas with the strongest positive or negative PC1 score displayed the strongest age-related increases or decreases, respectfully, in gradient position. However, there was noticeable variability in this relationship (e.g., some regions with a strong PC1 score displayed strong decreases; this would indicate that these regions became less preferentially connected to posterior areas across development). We tested an alternative hypothesis: that age-related changes occur along Cartesian thalamic planes (Oldham & Ball, 2023; Vogel et al., 2022). For PC1, we observed significant age-related changes in thalamic scores along medial-lateral (*r* = 0.32, *p* < .001), and anterior-posterior (*r* = −0.35, *p* < .001) axes (**Figure S6A-C**). Age related changes in PC2 and PC3 showed stronger correlations (compared to PC1) with their respective term template score/loadings, but inspection of these relationships shows that considerable heteroskedasticity in regional maturation of thalamic seeds and cortical vertices positioned along each axis (**Figure S5C-F**). These PC2 and PC3 thalamic age-related changes showed the strongest association with the ventral-dorsal plane (**Figure S6D-I**).

Finally, we examined if interruption to brain development in the third trimester affects macroscale patterns of thalamocortical connectivity. In total, n = 108 neonates included in this study were born preterm (born at less than 37 weeks gestation), a demographic in which abnormal thalamocortical connectivity has been previously observed (Ball et al., 2013, 2015; Toulmin et al., 2021). We selected a subset of scans where neonates born premature had been subsequently scanned at term equivalent age (n = 51; 25 females; median birth age [range] = 33^+1^ weeks [23^+4^-36^+6^]; median scan age [range] = 40^+4^ [37-45^+1^] weeks), and a set of controls who were born at term matched for age at scan and sex.

For each neonate, we calculated the total connectivity of each vertex to the thalamus (i.e., summed the connectivity across thalamic seeds for each vertex). Differences between preterm and term neonates thalamocortical cortical connections were measured by a two-tailed *t*-test, controlling for scan age and gender, with a Threshold-Free Cluster Enhancement correction (S. Smith & Nichols, 2009). Significant differences were detected by permutation testing (10,000 permutations) at a family-wise error rate of *p_FWER_* < .05, as implemented in Permutation Analysis of Linear Models (PALM) toolbox (Winkler et al., 2014).

We found significant differences in thalamocortical connectivity in preterm infants, compared to term controls, with decreased connectivity in frontal areas and occipital areas but moderate increases in parietal areas (**Figure 4**). After controlling for scan age and sex, the amount of variance explained by PC1 (mean±SD % variance explained; Term = 33.81±3.09%, Preterm = 33.78±2.29%; *F*(1,98) = 0.002, *p* = 0.997) and PC3 (Term = 7.38±0.62%, Preterm = 7.65±0.95%; *F*(1,98) = 0.002, *p* = 0.087) did not significantly differ between groups. The amount of variance explained by PC2 did differ between preterm and term neonates, although the difference was small (Term = 22.26±2.12%, Preterm = 21.04±2.13%; *F*(1,98) = 0.002, *p* = 0.001). In line with these results, preterm-term differences were only weakly correlated with PC loadings and age-related changes in PCs (**Figure S7**); indicating that the direction and magnitude of primary axes of thalamocortical connectivity were largely unaffected by preterm birth.

**Figure 4.**
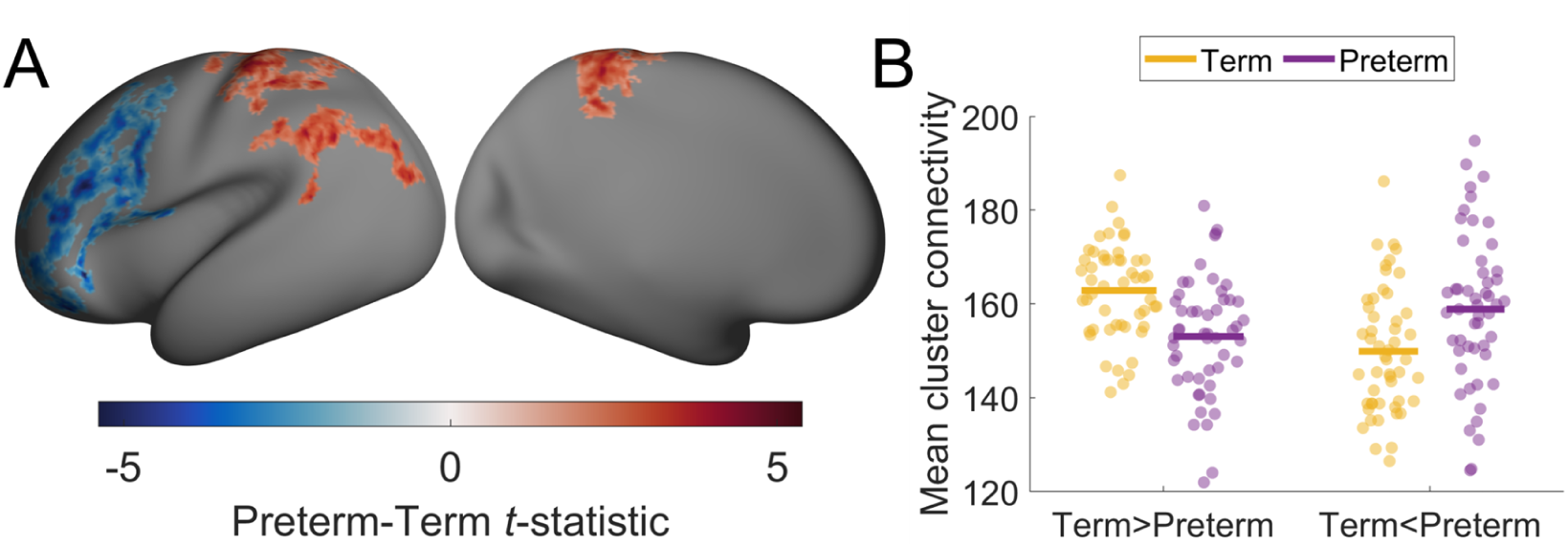
Significant differences in thalamocortical connectivity due to prematurity. (**A**) Differences between preterm and term neonate thalamocortical connectivity in the cortex. The *t* statistic is positive when preterm neonates had higher connectivity than term neonates and is negative when it was lower. Higher or lower values indicate the proportion of connectivity in that area was increased or decreased respectively when compared to what was expected in term neonates. Non-significant areas (*p_FWER_* < .05) are not coloured. (**B**) Mean connectivity of significant clusters. We took the mean connectivity of all vertices that showed a significant negative or positive difference in thalamocortical connectivity between preterm and term neonates (Term>Preterm corresponds to blue areas in **A**, while Term<Preterm corresponds to red areas). We then compared this mean connectivity between term (yellow) and preterm (purple) neonates. The line indicates the mean for each group/cluster.

## 4 Discussion

In this study, we describe macroscale patterns of thalamocortical connectivity using high-resolution tractography in a cohort of newborns infants. We observe a primary axis of variation orientated along an anterior/medial-to-posterior/lateral direction in the thalamus. This is consistent with previous findings in neonates (Wilson et al., 2023; Zheng et al., 2023), and adults (Oldham & Ball, 2023). The primary thalamic axis is associated a pattern of preferential connectivity to cortical areas that varies along a rostral-caudal direction, as observed in adults (Oldham & Ball, 2023). The topographic arrangement of thalamocortical fibres is observed across species (Brysch et al., 1990; Höhl-Abrahão & Creutzfeldt, 1991), and suggests a foundational principle of thalamic organisation. Our results suggest that this basic topographic pattern is present from the start of the third trimester. While our observations are limited to newborn infants, including those born preterm, recent work using fetal diffusion MRI has reported a similar topographical arrangement as early as the second trimester (Wilson et al., 2023).

Thalamocortical axons begin to gather in the subplate before innervating the cortex from mid-gestation (Kostović & Judaš, 2010) with their spatial distribution in this stage aligning with their eventual cortical targets (Molnár et al., 2012). As such, the primary topographical patterning of thalamocortical connections is likely established prior to the third trimester (Kostović & Judaš, 2010). Despite this, we observed significant refinement to thalamocortical connectivity strength along primary organisational axes between 30- and 45- weeks PMA. The changing strength of preferential connectivity to cortical areas are likely reflective of the maturation of existing thalamocortical fibres, potentially capturing processes that include both the removal of supporting radial glial structures and the onset of myelination (Machado-Rivas et al., 2021; Wilson et al., 2023).

We observed that age related changes in the pattern of connectivity associated with the primary thalamocortical component were most pronounced in medial-anterior, ventro-lateral, and ventral thalamic areas; with preferential connectivity to cortical association areas. Previous studies of the dHCP cohort have reported that the microstructure of thalamic subdivisions develop along a lateral-to-medial temporal axis, whilst thalamic connections to the cortex develop along a posterior-anterior axis (Zheng et al., 2023). Our results unify these observations by demonstrating that observed thalamic and cortical changes reflect refinements to the underpinning topographic pattern of thalamocortical connections. Furthermore, cortical association areas showed the greatest changes in thalamocortical connectivity during the third trimester, which aligns with broader cortical developmental patterns of delayed maturation in association and frontal areas (Tau & Peterson, 2010). Thalamocortical innervations to cortical association areas occur later in development (Altman & Bayer, 1979, 1988a, 1988b, 1988c; Chalfin et al., 2007; Finlay & Uchiyama, 2020), matching the pattern of thalamocortical development observed in our results. Evidence suggests there is considerable interplay between thalamic and cortical development (Antón-Bolaños et al., 2018; Finlay & Uchiyama, 2015, 2020; Park et al., 2024). Thalamocortical connectivity contributes to the arealisation of the cortex (Cadwell et al., 2019; Finlay & Uchiyama, 2015, 2020; Molnár & Kwan, 2024), and are involved in shaping the laminar organisation of the cortex (Dehay et al., 2001; Molnár & Kwan, 2024; Pouchelon et al., 2014; Sato et al., 2022), regulating cortical progenitors (Gerstmann et al., 2015), connectivity patterns (Finlay, 1991; Finlay & Uchiyama, 2020), and the initialisation of functional dynamics (Antón-Bolaños et al., 2018; Martini et al., 2021). According to the “handshake hypothesis”, ascending thalamic and descending cortical fibres meet in the subplate and reciprocally guide the tracts to their respective cortical and thalamic targets (Molnár et al., 2012; Molnár & Blakemore, 1995), further suggesting interactions between cortical and thalamic development are vital to shaping both structures. Our observations of age-related changes in thalamocortical connectivity fit with these wider findings that underscore the interdependence between thalamic and cortical development to shape the maturation and functional organization of the developing brain. As thalamocortical connectivity is related to multiple facets of cortical organisation (Oldham & Ball, 2023), and is considered to constrain how this organisation is established (Park et al., 2024), further investigation of the relationship between thalamocortical connectivity and major cortical organisational properties (e.g., integration/segregation, intrinsic timescales, cortical thickness) is warranted to more deeply understand the role of these connections in neurodevelopment.

We note that changes in the strength of PC scores and loadings with age were closely aligned with Cartesian anterior-posterior, medial-lateral, and inferior-superior axes. Spatial axes constitute a critical foundation for early brain development, corresponding to the direction of early molecular gradients (Vogel et al., 2022). Thalamic microstructure varies along cardinal planes (Altman & Bayer, 1979; Nakagawa, 2019; Scholpp & Lumsden, 2010; Wong et al., 2018) and neurogenesis progresses from the lateral to medial thalamus (Altman & Bayer, 1979; Nakagawa, 2019; Wong et al., 2018) with thalamic subdivisions also emerging along this same axis (Zheng et al., 2023), and myelination of white matter tracts occurring along an anterior-posterior direction (Abe et al., 2004; Counsell et al., 2002; Machado-Rivas et al., 2021). Therefore, the orientation of the primary thalamocortical connectivity components defined here may arise from the intersection of different developmental gradients. The temporal interaction of these gradients could account for the observed heterogeneity in age-related changes across thalamocortical axes.

Linear and nonlinear decomposition methods have become increasingly common in the neuroimaging literature to generate low-dimensional representations of complex imaging data (Margulies et al., 2016; Oldehinkel et al., 2023; Paquola et al., 2020; Vos De Wael et al., 2020). However, recent studies have highlighted potential biases in PCA decompositions of spatially autocorrelated data (Novembre & Stephens, 2008; Shinn, 2023; Watson & Andrews, 2023), whereby PCs can resolve into distinctive sinusoidal patterns that do not necessarily reflect the true structure of the underlying data. To avoid interpreting ‘phantom’ oscillations, we turn to converging lines of evidence from imaging (Lambert et al., 2017; Oldham & Ball, 2023; Pang et al., 2023; Yang et al., 2020; Zheng et al., 2023), histological/cellular data (Roy et al., 2022), animal tract tracing (Brysch et al., 1990; Höhl-Abrahão & Creutzfeldt, 1991), and transcriptomics (Phillips et al., 2019; Vogel et al., 2022) that support the organisation of thalamic neurobiology along major cardinal planes, like those we observe in the current study. These convergent findings using different modalities indicate the observed axis are not merely a statistical artefact. There are distinctive spatiotemporal dynamics of molecular expression during thalamic development (Kim et al., 2023), which likely drive the formation of continuous axes of neurobiological features (Le Dréau & Martí, 2012; Sansom & Livesey, 2009). Determining how various organisational axes of different brain systems are aligned and functionally related; how common or distinct mechanisms may shape their emergence; and how these patterns are related to cognition, behaviour, sensation, and symptomology; is key to understanding thalamocortical organisation and development.

The nuclear organisation of the thalamus is extremely well characterised (Jones, 2007; Phillips et al., 2019; Roy et al., 2022), and distinct developmental processes are associated with different nuclei (Huang et al., 2024; Nakagawa, 2019). The presence of continuous developmental gradients and axis of connectivity should be considered a complementary organisational principle. For example, the core-matrix (Jones, 1998) and higher-first order theories of thalamic organisation (Guillery & Sherman, 2002; Sherman, 2016; Sherman & Guillery, 2002) propose that nuclei have both diffuse and focal cortical targets (with nuclei varying in their ratio of these connection types). These diffuse connections, which extend across multiple cortical areas (Jones, 1998), potentially correspond to the principal components defined herein, rather than the more areal specific, targeted connectivity that defines focal/core connections (Jones, 1998) as the extracted connectivity components smoothly vary across the cortex without respect to areal boundaries. In addition, diffuse and specific thalamocortical connections may emerge due to distinctive developmental events. For example, matrix axons have been suggested to invade the cortex prior to core axons (Clascá et al., 2012), or alternatively that core/first-order connections develop from matrix/higher-order connections (Lo Giudice et al., 2024). By definition, PCA extracts patterns which explain the most variance in the data, therefore the first few components are expected to define widespread connectivity patterns. This does not preclude the existence of focal connectivity patterns, which are a highly important aspect of thalamic organisation and function (Bosch-Bouju et al., 2013; Fama & Sullivan, 2015; Guillery & Sherman, 2002; Jones, 2007), but these are likely to explain less variance in the data as such connectivity will only apply to a select subset of thalamic seeds. Establishing methods which can disentangle the focal and diffuse connectivity patterns of the thalamus is important to allow more detailed exploration of thalamic organisation and its development (Howell et al., 2024).

We also observed significant differences in thalamocortical connectivity between preterm and term-born neonates. At term-equivalent age, preterm infants had a lower proportion of thalamic connections to frontal and occipital areas compared to term-born peers, with some parietal areas showing a higher proportion. These areas affected by prematurity are similar to what previous studies have reported (Ball et al., 2012, 2013, 2015; Batalle et al., 2017; Jakab et al., 2020; Sa De Almeida et al., 2021), although the abnormalities we observed were not as widespread or pronounced. This could be because in previous studies preterm neonates had a greater degree of prematurity than the cohort considered in this study, and neurodevelopmental abnormalities scale with the severity of preterm birth (Drommelschmidt et al., 2024). Premature birth disrupts normal developmental processes that shape brain connectivity through inflammation, infection, and/or perinatal hypoxia (Deng, 2010; Volpe, 2009). In particular, preterm birth is linked to subplate damage, a structure critical to enabling a variety of maturational processes like synaptogenesis and axonal guidance (Volpe, 2009). Therefore, damage to it may impair the proper establishment of thalamocortical connectivity. Whilst we observed impaired connectivity in the preterm neonates of this study, these impairments were only weakly correlated with the major axes of thalamocortical connectivity, indicating that prematurity did not fundamentally change the core organisational topography of thalamocortical connections. As the topographical organisation of thalamic connections is established early in gestation (Kostović et al., 2019; Kostović & Judaš, 2010) and diffuse connections form potentially prior to functionally specialised, focal connections (Clascá et al., 2012; Lo Giudice et al., 2024), the adverse effects of preterm birth may be more evident in focal thalamocortical connections, rather than macroscale, diffuse organisational patterns but further characterisation of focal and diffuse thalamocortical connectivity during early neurodevelopment will be required to further advance this hypothesis.

Our study has several limitations which are important to note. First, we used MRI scans of preterm infants to investigate thalamocortical development prior to the time of normal birth. Whilst prematurity does alter the developmental trajectory of structural thalamocortical connectivity (Ball et al., 2012, 2013; Batalle et al., 2017; Sa De Almeida et al., 2021), the major developmental trends - such as would be captured by the first few components - are likely to be more pronounced than such differences this would introduce (Zhao et al., 2019; Zheng et al., 2023). This is supported by our findings’ showing prematurity was only weakly related to thalamocortical gradient patterning. Secondly, we only focused the left hemisphere in our analysis, as the majority of thalamocortical connections are ipsilateral (Dermon & Barbas, 1994). However, for a complete representation of thalamocortical connectivity, future studies should endeavour to measure connections across both hemispheres. A more accurate characterisation of thalamocortical connectivity may also be achieved by using alternative measures to define white matter connections. Using measures of connectivity aimed at more explicitly modelling the biological properties of axonal fibres (Raffelt et al., 2017; R. E. Smith et al., 2022; F. Zhang et al., 2022; H. Zhang et al., 2012) may yield deeper insights into the maturational patterns of thalamocortical connectivity. Thirdly, conducting tractography from the deep thalamus is challenging which is compounded by using a neonatal cohort. Neonates are more susceptible to head motion, which is disruptive to diffusion sequences (Heemskerk et al., 2013; Pannek et al., 2012), which can be highly difficult to correct for and so residual motion artefacts may still be present in our data. We used data processing with optimised pipelines to migrate these issues (Bastiani et al., 2019) and employed strict quality control to minimise any impact to our results. Our thalamocortical findings are consistent with histological and animal studies (Brysch et al., 1990; Höhl-Abrahão & Creutzfeldt, 1991), and are in agreement with thalamocortical connectivity observed in adult imaging data (Howell et al., 2024; Oldham & Ball, 2023), and other early life studies (Wilson et al., 2023; Zheng et al., 2023), adding further confidence our results are not adversely affected by motion induced distortions.

In summary, this study investigated the development of thalamocortical connectivity in the perinatal period. We find that a primary thalamocortical axis, describing changing patterns of cortical connectivity along an anterior/medial-to-posterior/lateral orientation in the thalamus is established by 30 weeks gestation. Changes to this axis prior to the time of normal birth is largely driven by maturation of connections between the thalamus and associative cortical areas. Finally, thalamocortical connectivity differences due to prematurity were only weakly related to thalamocortical axes, suggesting the conservation of these major organisational features following preterm birth.

## Acknowledgements

The authors would like to thank Ashlea Segal for providing technical assistance for the analysis. This research was supported by an NHMRC Investigator Grant (1194497 to G.B.), the Murdoch Children’s Research Institute, the Royal Children’s Hospital, Department of Paediatrics, The University of Melbourne and the Victorian Government’s Operational Infrastructure Support Program. The project was generously supported by The Royal Children’s Hospital Foundation devoted to raising funds for research at The Royal Children’s Hospital. Data were provided by the developing Human Connectome Project, KCL-Imperial-Oxford Consortium funded by the European Research Council under the European Union Seventh Framework Programme (FP/2007-2013) / ERC Grant Agreement no. [319456]. We are grateful to the families who generously supported this trial.

## Data and Code Availability

Neuroimaging data for the developing Human Connectome Project are available on the NIMH Data Archive. Instructions on how to access are available here: https://biomedia.github.io/dHCP-release-notes/. Processed data is available on Zenodo (https://zenodo.org/doi/10.5281/zenodo.11059161) and code is available https://github.com/StuartJO/ThalamicDevGrad.

## Author Contributions

Stuart Oldham: Conceptualization, Methodology, Formal analysis, Resources, Data Curation, Writing - Original Draft, Writing - Review & Editing, Visualization. Sina Mansour L: Methodology, Writing - Review & Editing. Gareth Ball: Conceptualization, Supervision, Project administration.

## Supplementary material

**Figure S1.**
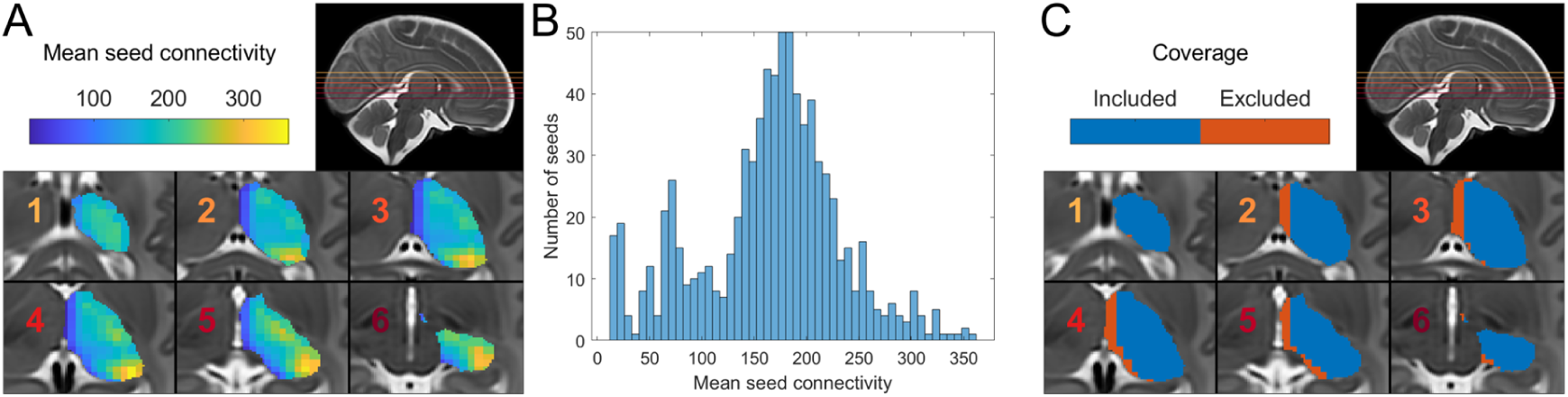
Distribution of non-normalised connectivity values for each seed. **(A)** The non-normalised connectivity from each thalamic seed to the cortex (averaged across all neonates). (**B**) Histogram of seed connectivity values. (**C**) Final selected seed coverage of the thalamus. Orange areas, whilst covered by the thalamic mask, were excluded as they showed noticeably weaker connectivity than other thalamic areas.

**Figure S2.**
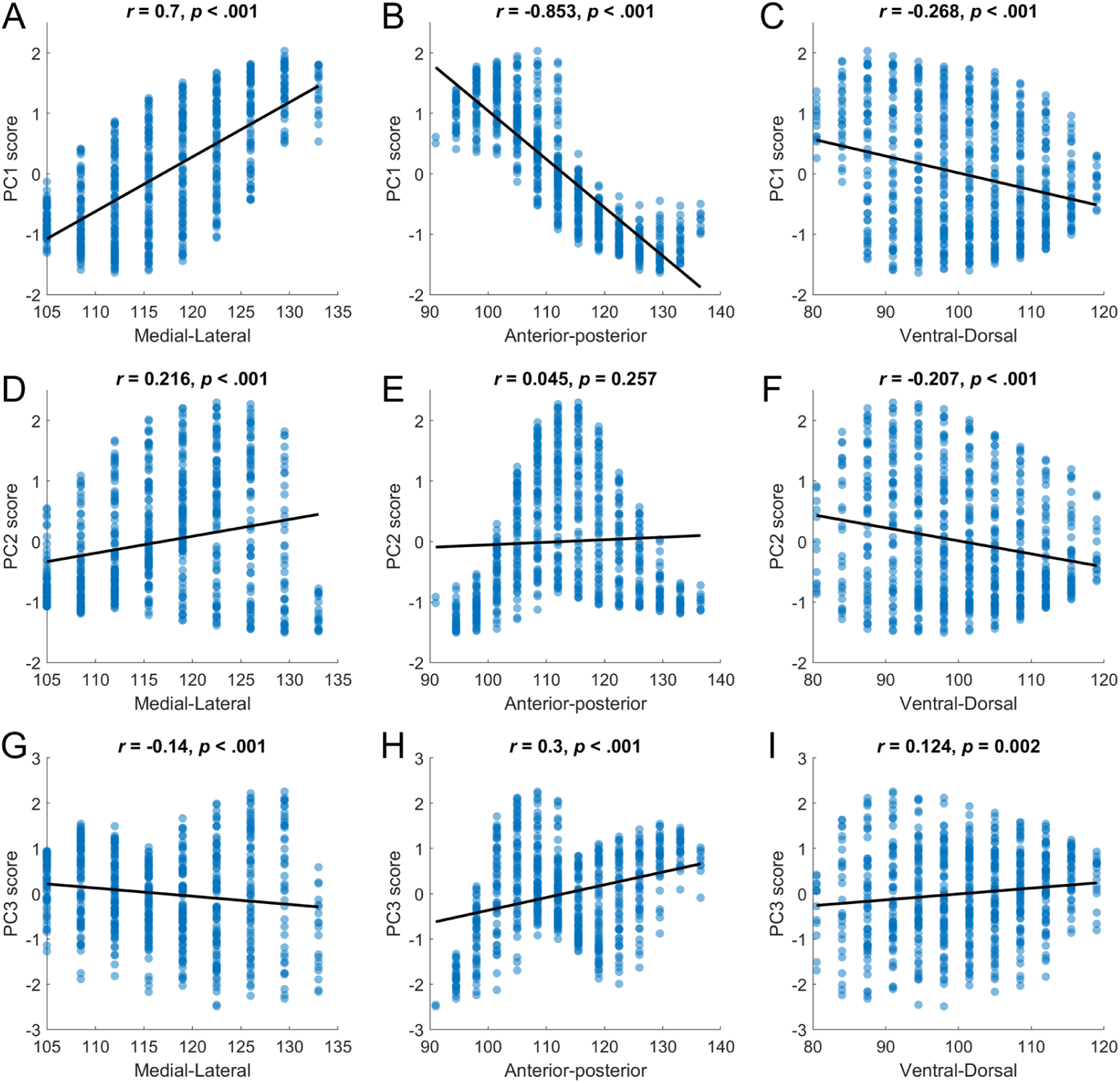
Correlations between seed PC1-PC3 score and cartesian axes position. **(A)** PC1 score with Medial-lateral axis (*x*-axis voxel coordinate). **(B)** PC1 score with anterior-posterior axis (*y*-axis voxel coordinate). **(C)** PC1 score with dorsal-ventral axis (*z*-axis voxel coordinate). **(D)** PC2 score with Medial-lateral axis (*x*-axis voxel coordinate). **(E)** PC2 score with anterior-posterior axis (*y*-axis voxel coordinate). **(F)** PC2 score with dorsal-ventral axis (*z*-axis voxel coordinate). **(G)** PC3 score with Medial-lateral axis (*x*-axis voxel coordinate). **(H)** PC3 score with anterior-posterior axis (*y*-axis voxel coordinate). **(I)** PC3 score with dorsal-ventral axis (*z*- axis voxel coordinate). The black line indicates the (linear) line of best fit.

**Figure S3.**
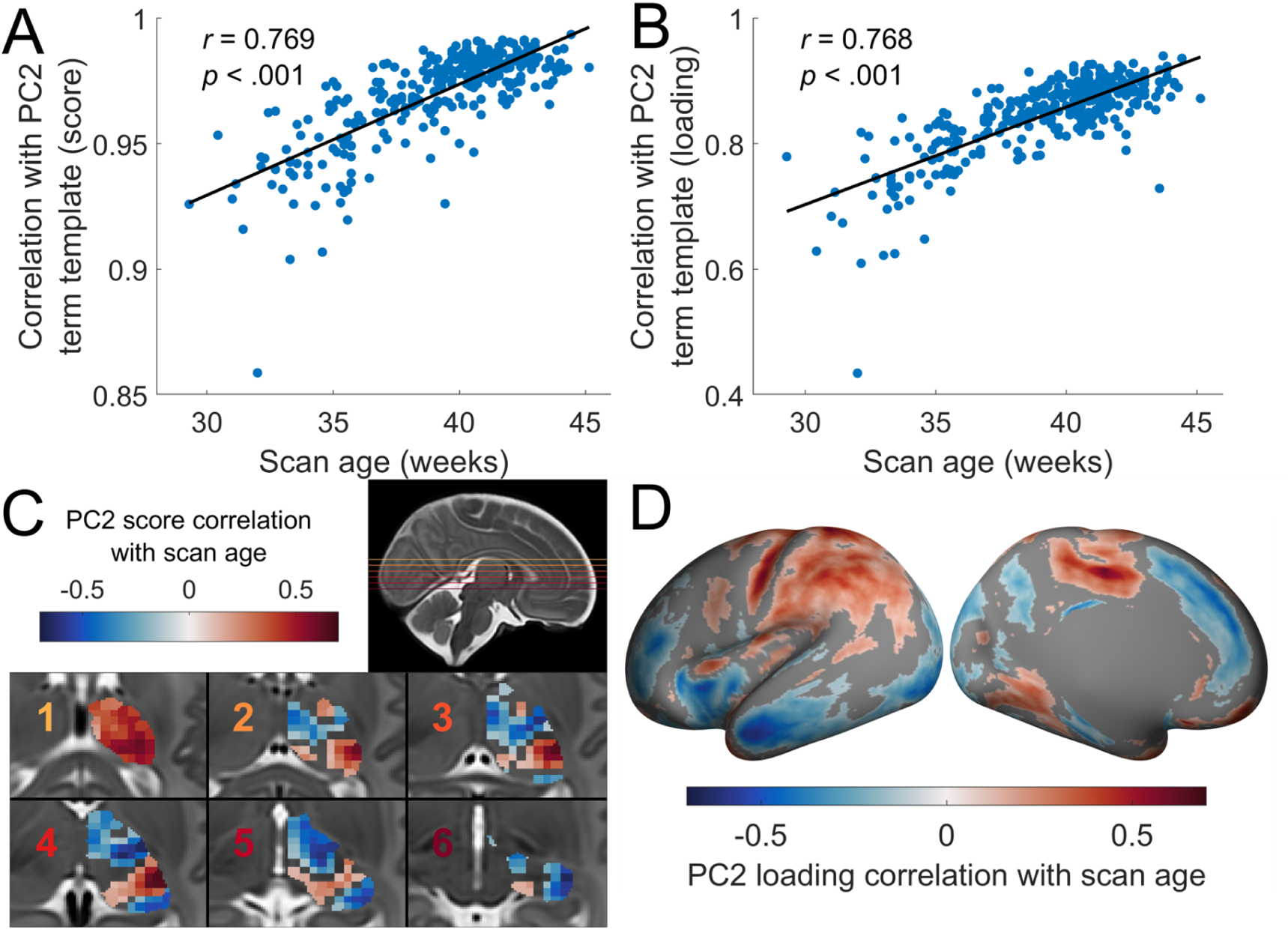
Age-related changes in the secondary thalamocortical axis. (**A**) Scatter plot of the correlation between individual and template PC2 scores and individual scan age. (**B**) Scatter plot of the correlation between individual and template PC2 loading and individual scan age. (**C**) Correlation between individual PC2 scores and scan age for each thalamic seed (voxels are coloured according to the value of the nearest seed; non-significant areas are not coloured; *p_FDR_* < 0.05). (**D**) Correlation between individual PC2 loading and scan age for each cortical vertex (non-significant areas are not coloured; *p_FDR_* < 0.05).

**Figure S4.**
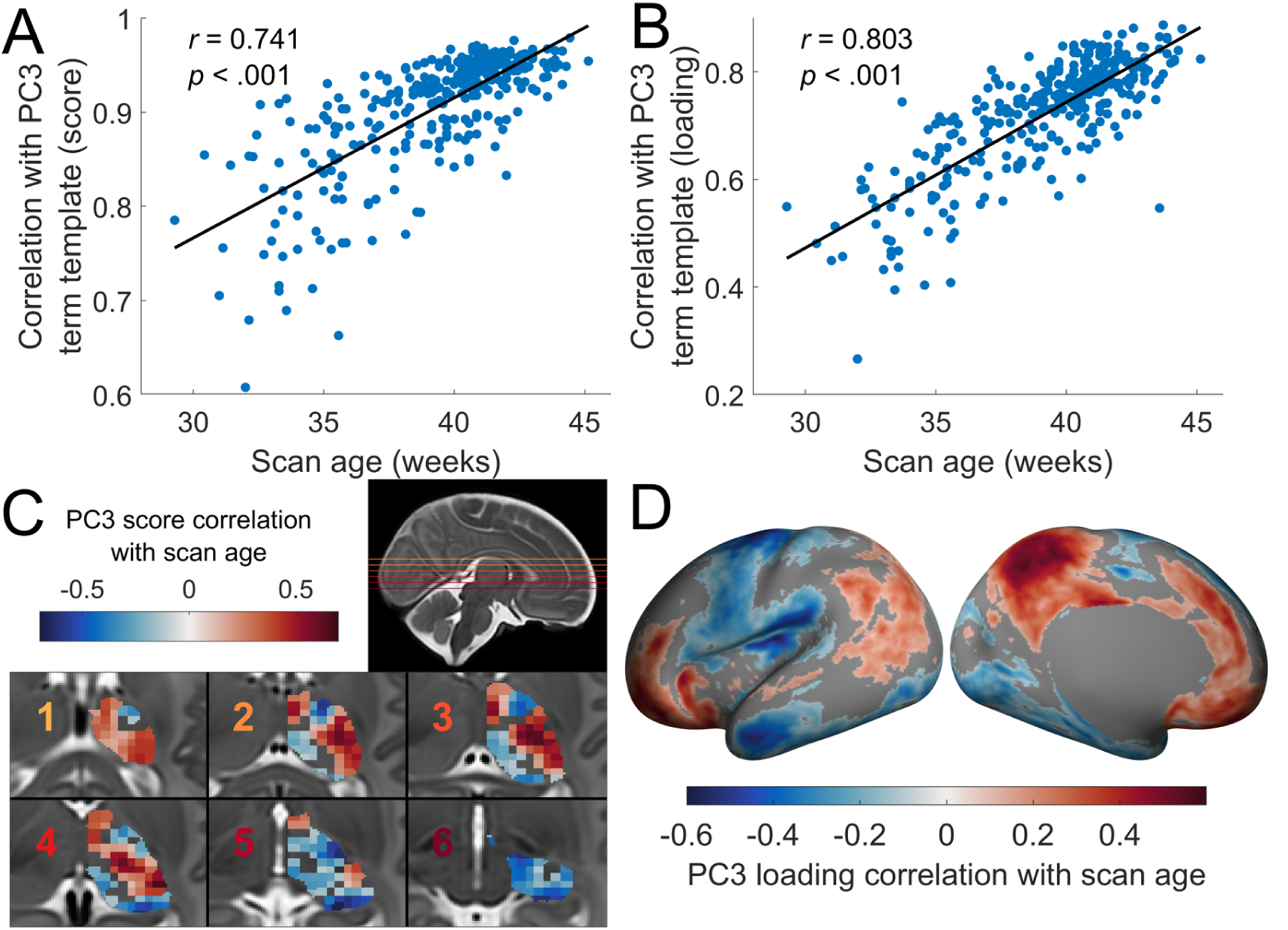
Age-related changes in the tertiary thalamocortical axis. (**A**) Scatter plot of the correlation between individual and template PC3 scores and individual scan age. (**B**) Scatter plot of the correlation between individual and template PC3 loading and individual scan age. (**C**) Correlation between individual PC3 scores and scan age for each thalamic seed (voxels are coloured according to the value of the nearest seed; non-significant areas are not coloured; *p_FDR_* < 0.05). (**D**) Correlation between individual PC3 loading and scan age for each cortical vertex (non-significant areas are not coloured; *p_FDR_* < 0.05).

**Figure S5.**
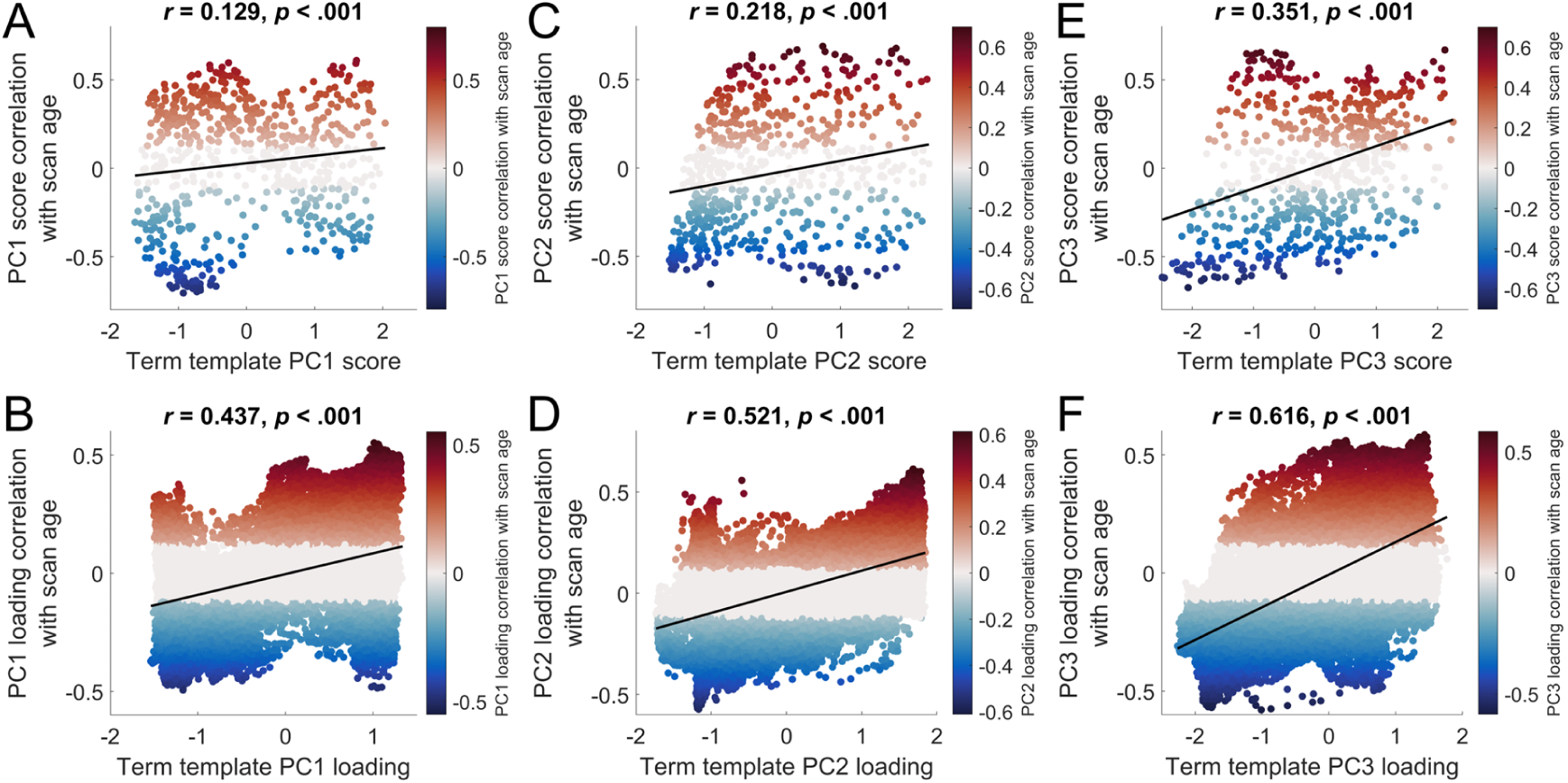
Relationship between age-related gradient changes and gradient position. (**A**) Relationship between the term template PC1 score and PC1 score age-related changes. (**B**) Relationship between the term template PC1 loading and PC1 loading age-related changes. (**C**) Relationship between the term template PC2 score and PC2 score age-related changes. (**D**) Relationship between the term template PC2 loading and PC2 loading age-related changes. (**E**) Relationship between the term template PC3 score and PC3 score age-related changes. (**F**) Relationship between the term template PC3 loading and PC3 loading age-related changes. The black line indicates the line of best fit.

**Figure S6.**
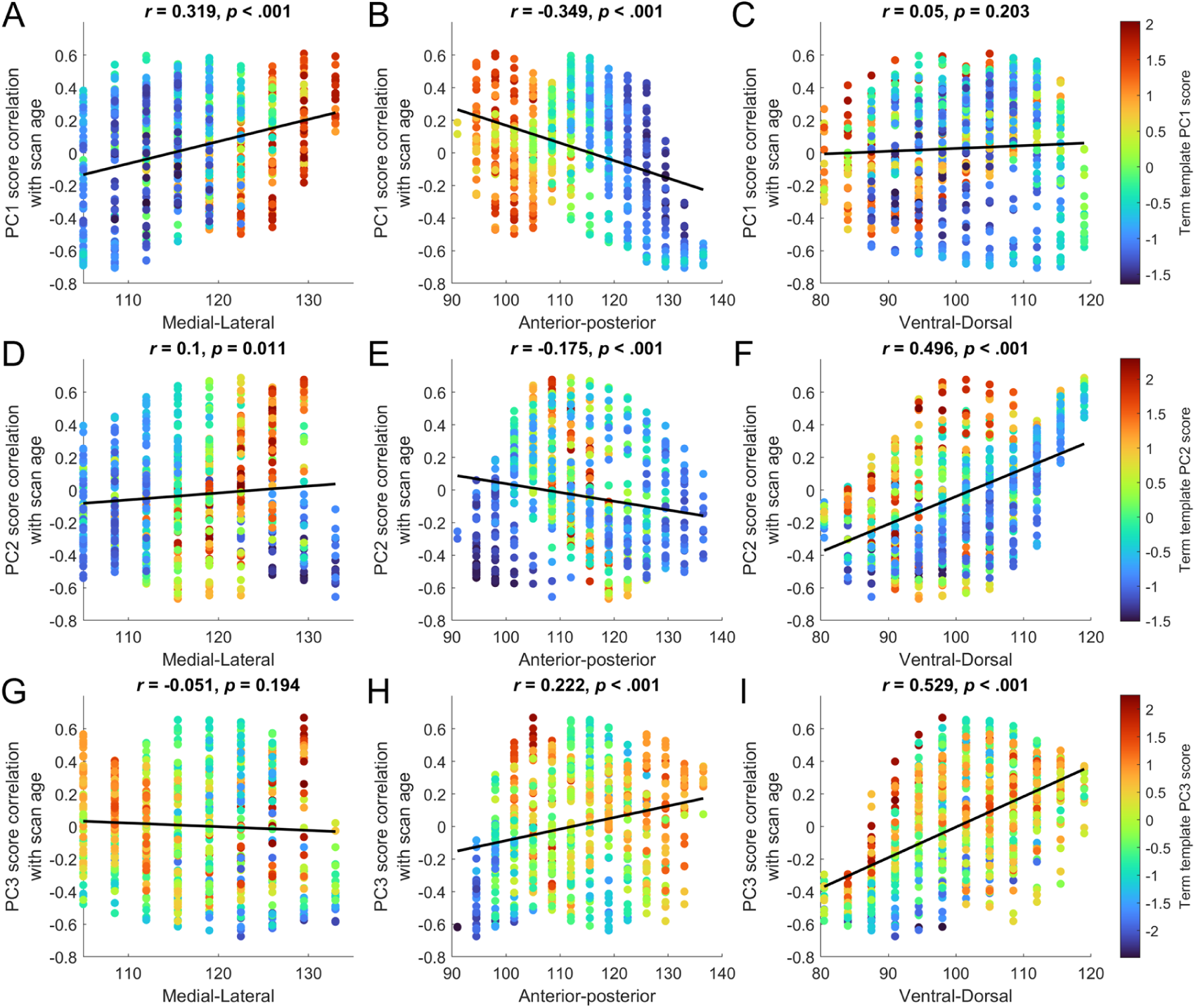
Correlations between age-related changes in PC score and cartesian axes position. **(A)** Relationship between age-related changes in PC1 scores and with medial-lateral axis position (*x*-axis voxel coordinate). **(B)** Relationship between age-related changes in PC1 scores and with anterior-posterior axis (*y*-axis voxel coordinate). **(C)** Relationship between age-related changes in PC1 scores and with dorsal-ventral axis (*z*-axis voxel coordinate). **(D)** Relationship between age-related changes in PC2 scores and with medial-lateral axis position (*x*-axis voxel coordinate). **(E)** Relationship between age-related changes in PC2 scores and with (*y*-axis voxel coordinate). **(F)** Relationship between age-related changes in PC2 scores and with (*z*-axis voxel coordinate). **(G)** Relationship between age-related changes in PC3 scores and with medial-lateral axis position (*x*-axis voxel coordinate). **(H)** Relationship between age-related changes in PC3 scores and with anterior-posterior axis (*y*-axis voxel coordinate). **(I)** Relationship between age-related changes in PC3 scores and with dorsal-ventral axis (*z*-axis voxel coordinate). The black line indicates the line of best fit.

**Figure S7.**
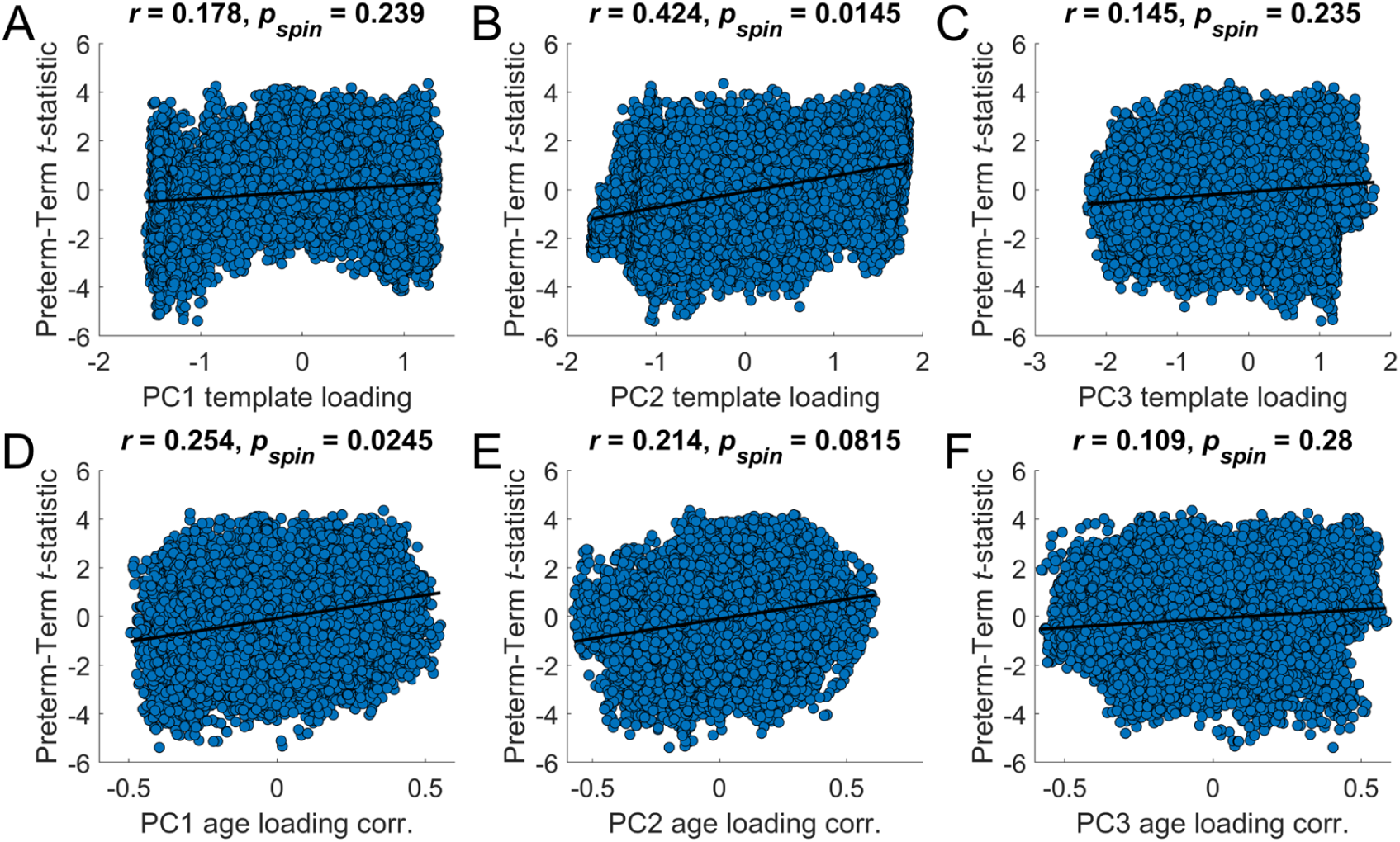
Relationship between cortical thalamocortical connectivity preterm-term differences and gradient position/age-related changes. (**A**) Relationship between preterm-term thalamic connectivity difference and PC1 loading. (**B**) Relationship between preterm-term thalamic connectivity difference and PC2 loading. (**C**) Relationship between preterm-term thalamic connectivity difference and PC3 loading. (**D**) Relationship between preterm-term thalamic connectivity difference and PC1 loading age-related changes. (**E**) Relationship between preterm-term thalamic connectivity difference and PC2 loading age-related changes. (**F)** Relationship between preterm-term thalamic connectivity difference and PC3 loading age-related changes. The black line indicates the line of best fit. Significance was determined using a spin test (*p*_*spin*_ < .05).

